# The role of mitochondrial energetics in the origin and diversification of eukaryotes

**DOI:** 10.1101/2021.10.23.465364

**Authors:** Paul E. Schavemaker, Sergio A. Muñoz-Gómez

**Affiliations:** Center for Mechanisms of Evolution, The Biodesign Institute, School of Life Sciences, Arizona State University, 727 E. Tyler St. Tempe, AZ 85281-5001, U.S.A; Unité d’Ecologie, Systématique et Evolution, Université Paris-Saclay, Orsay, France

**Author notes:** Equal contribution.

**Keywords:** Eukaryogenesis, energy, complexity, genome size, cell volume

## Abstract

The origin of eukaryotic cell size and complexity is thought by some to have required an energy excess provided by mitochondria, whereas others claim that mitochondria provide no energetic boost to eukaryotes. Recent observations show that energy demand scales continuously and linearly with cell volume across both prokaryotes and eukaryotes, and thus suggest that eukaryotes do not have an increased energetic capacity over prokaryotes. However, amounts of respiratory membranes and ATP synthases scale super-linearly with cell surface area. Furthermore, the energetic consequences of the contrasting genomic designs between prokaryotes and eukaryotes have yet to be precisely quantified. Here, we investigated (1) potential factors that affect the cell volumes at which prokaryotes become surface area-constrained, and (2) the amount of energy that is divested to increasing amounts of DNA due to the contrasting genomic designs of prokaryotes and eukaryotes. Our analyses suggest that prokaryotes are not necessarily constrained by their cell surfaces at cell volumes of 10^0^‒10^3^ μm^3^, and that the genomic design of eukaryotes is only slightly advantageous at genomes sizes of 10^6^‒10^7^ bp. This suggests that eukaryotes may have first evolved without the need for mitochondria as these ranges hypothetically encompass the Last Eukaryote Common Ancestor and its proto-eukaryotic ancestors. However, our analyses also show that increasingly larger and fast-dividing prokaryotes would have a shortage of surface area devoted to respiration and would disproportionally divest more energy to DNA synthesis at larger genome sizes. We thus argue that, even though mitochondria may not have been required by the first eukaryotes, the successful diversification of eukaryotes into larger and more active cells was ultimately contingent upon the origin of mitochondria.

**Significance:** There has been a lot of theorizing about the evolution of eukaryotes from prokaryotes, but no consensus seems to be on the horizon. Our quantitative analyses on the required amount of respiratory membrane, and the amount of energy diverted to DNA synthesis, by both prokaryotes and eukaryotes, suggest that mitochondria provided rather small advantages to the first eukaryotes, but were advantageous for the macro-evolutionary diversification of eukaryotes. This conclusion provides a middle road in the debate between those that claim that the origin of eukaryotes required a massive energy boost provided by mitochondria, and those that argue that the origin of mitochondria did not represent a quantum leap in energetic advantages to eukaryotes.

## Introduction

The transition from prokaryotic to eukaryotic cells is often thought to be the greatest transition in the history of life (1). This is because this is the largest gap, or discontinuity, in organismal structure or organization across the tree of life: eukaryotic cells are structurally much more complex, and on average, also larger in volume than prokaryotic cells (2). Many authors have thus attempted to explain how eukaryotes evolved from prokaryotes (3–9). However, much debate and speculation persist about the processes that gave rise to the first eukaryote (10, 11, 3, 12).

To explain the apparent large gap or gulf in anatomical complexity between prokaryotes and eukaryotes, the energetic hypothesis for eukaryote genome complexity suggests that there is also a deep energetic divide between these two grades of organization ((3) and see (8, 13–15) for precursors). Lane and Martin claim that eukaryotes have, on average, ~200,000 times more ‘energy per gene’ than prokaryotes (3). Such a drastic energetic difference is supposedly caused by two major advantages conferred by mitochondria upon eukaryotes (3, 16–18). The first one is the internalization and expansion of respiratory membranes within mitochondria which released eukaryotes of surface-area constraints. The second one is the evolution of highly reduced and specialized mitochondrial genomes which conferred a genomic asymmetry upon eukaryotes. Unlike prokaryotes, which have a single genome that scales up in number proportionally with cell volume, eukaryotes have a single large nuclear genome whose copy number can remain constant and numerous much smaller mitochondrial genomes that scale up in number with cell volume. The combination of these two advantages, according to Lane and Martin, allowed a drastic increase in the energy available per gene expressed in eukaryotes relative to prokaryotes (3, 16–18). One possible interpretation of this hypothesis predicts a jump in energetic capacity that separates eukaryotes from prokaryotes (Fig. 1A). Mitochondria are the cause of these massive energetic differences, Lane and Martin argue, and were thus a pre-requisite for the evolution of the eukaryotic complexity (3, 18).

**Figure 1.**
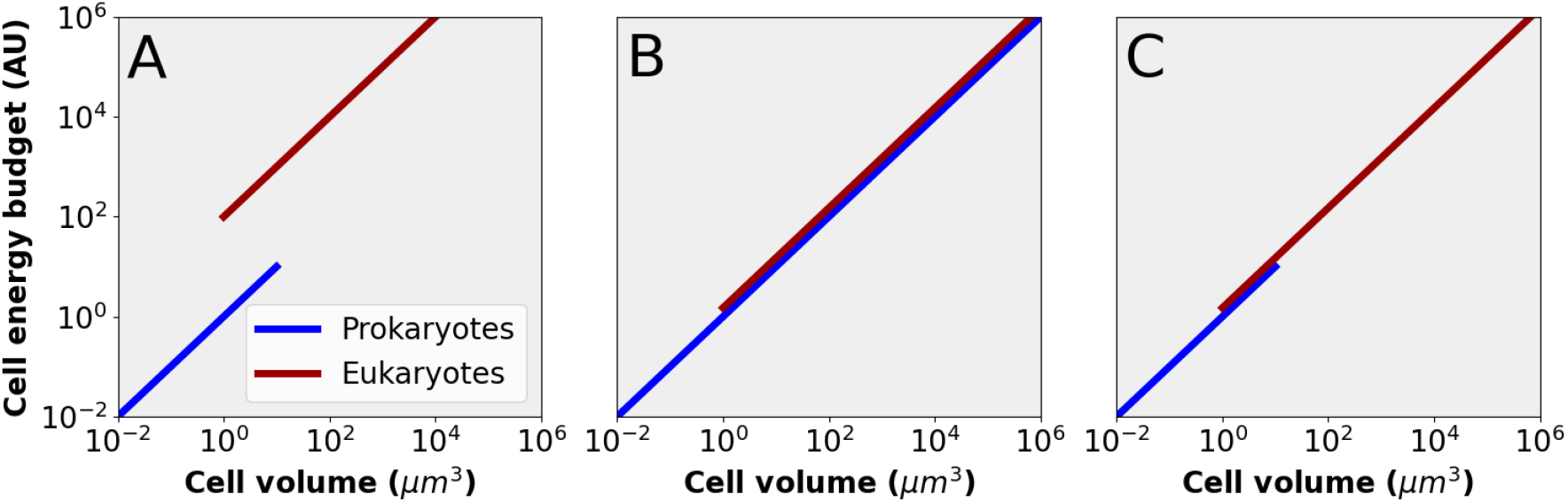
Three different possibilities for the energetic scaling across cell volume for prokaryotes and eukaryotes. **A**. A discontinuity in the scaling of cell energy with cell volume between prokaryotes and eukaryotes, where the latter exhibit a higher energetic capacity or energy density due to mitochondria. **B**. The hypothetical situation in the absence of surface constraints to prokaryotic cell volume, with the energetic capacity of prokaryotes accompanying that of eukaryotes over the full volume range. **C**. Continuous scaling of cell energy with cell volume over the prokaryote-eukaryote divide, based on data presented by Lynch and Marinov (10), and Chiyomaru and Takemoto (19). Unlike in B, the cell volume of prokaryotes is constrained. This constraint may be caused by the lack of a cytoskeleton, endomembrane system, or mitochondrion-based respiration.

Some authors have expressed skepticism about the energetic hypothesis for the origin of eukaryotic complexity (10, 19–24). The notion that the evolution of cell complexity requires an increased energy supply has been dismissed as having no evolutionary basis (22, 24), and the concept of ‘energy per gene’ has been criticized as evolutionarily meaningless (23, 25). Recently, Lynch and Marinov found a continuous energetic scaling across prokaryotes and eukaryotes (10), and similar results have been presented by Chiyomaru and Takemoto (19). This suggests that there is no energetic gap (or shift in energetic capacity) between prokaryotes and eukaryotes, as the amount of energy available to a cell is directly proportional to its volume regardless of whether the cell is prokaryotic or eukaryotic. Based on this, Lynch and Marinov argue that mitochondria do not provide eukaryotes with a higher energetic capacity and imply that prokaryotes are energetically unconstrained by their cell surfaces (Fig. 1B). Moreover, Lynch and Marinov showed that the number of ATP synthases scales continuously across prokaryotes and eukaryotes, and argued that the increase in surface area provided by mitochondria is not particularly large when compared to that available at the cytoplasmic membrane (21). However, their data also show that the amount of mitochondrial membrane and the number of ATP synthases scale super-linearly with the cell surface area (21). This suggests, in contrast to Lynch and Marinov (10, 21), that prokaryotes might be constrained by their cell surfaces at larger volumes, and that mitochondria may allow eukaryotes to scale up in cell volume without a shortage of respiratory membranes (Fig. 1C). Furthermore, the energetic consequence of the contrasting genomic designs between prokaryotes and eukaryotes, first emphasized by Lane and Martin (3, 16) but ignored by others, remains to be explored.

To explore the potential energetic benefits that mitochondria bestowed upon eukaryotes, our goal here has been to carefully dissect major differences between mitochondrion-less and mitochondrion-bearing cells (i.e., prokaryotes and eukaryotes, respectively) in light of the recent scaling laws devised by Lynch and Marinov (10, 21). To do so, we (1) explored potential factors (cell shape, cell division time, and maximum fraction of respiratory membrane) that affect the cell volumes at which anatomically simple cells become surface area-constrained, and (2) investigated the decrease in energy budget that is associated with the contrasting genomic designs exhibited by mitochondrion-less and mitochondrion-bearing cells across a wide range of cell volumes. We discuss our observations in the context of the prokaryote-eukaryote divide and the origin and diversification of eukaryotes.

## Results

### The cell volume, genome size, and gene number distributions of prokaryotic and eukaryotic cells

In this manuscript, we use theoretical models to assess the respiratory membrane requirements and DNA investments of mitochondrion-less and mitochondrion-bearing cells. These models might help explain, from an energetic point of view, the differences observed between modern prokaryotes and eukaryotes and thus inform our discussions of the prokaryote-eukaryote transition. We start by presenting the distributions of cell volume, genome size and gene number from a comprehensive survey of phylogenetically disparate prokaryotes and eukaryotes (Fig. 2; Dataset S1).

**Figure 2.**
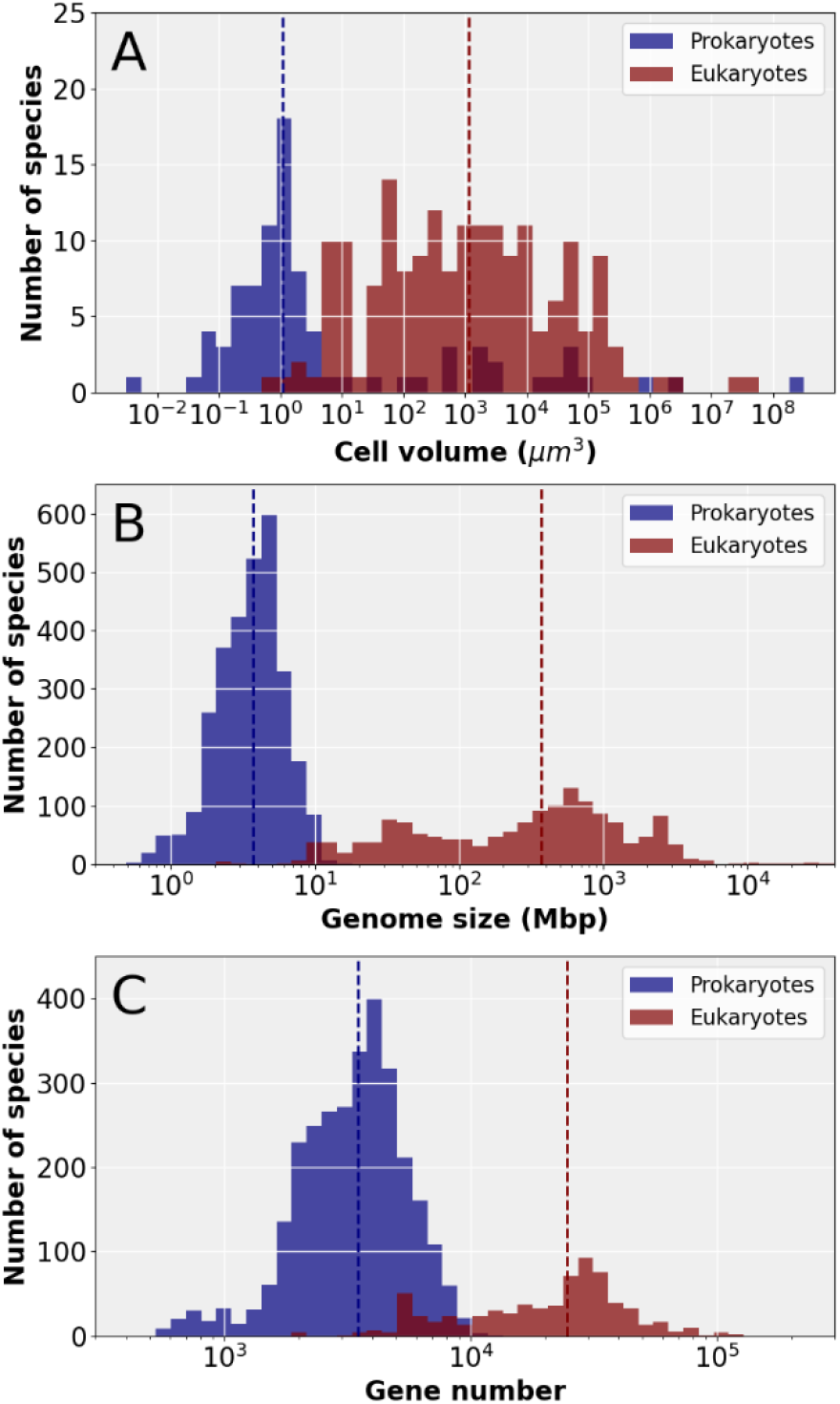
Cell volumes, genome sizes, and gene numbers for prokaryotes and eukaryotes. Cell volumes for diverse eukaryotes were obtained from (10) and additional data were added from several sources (see Dataset S1). Genome sizes and gene numbers were acquired from NCBI GenBank and manually curated to remove outliers due to gene mis annotations. The vertical dashed lines show medians. Total cell volumes, instead of active cytoplasmic volume, were used for giant prokaryotes (>10^2^ μm^3^).

The cell volume distributions of prokaryotes and eukaryotes point at two main conclusions. First, the ranges for each grade of organization (~10^−2^‒10^2^ μm^3^ for prokaryotes and ~10^0^‒10^8^ μm^3^ for eukaryotes) do not overlap for the most part: their medians (vertical dashed lines in Fig. 2) largely fall outside of each other’s distributions (Fig. 2A). This is most obvious when giant bacteria like *Beggiatoa* spp. and *Thiomargarita namibiensis* are excluded (Fig. 2A; Dataset S1). Giant bacteria reach absolute volumes >10^6^ μm^3^ but these are mostly inert as they contain a large central vacuole or numerous intracellular inclusions made of sulfur or calcium carbonate (some exceptions are large cyanobacterial cells; Dataset S1). Thus, most prokaryotes are smaller than most eukaryotes. Second, prokaryotes and eukaryotes mostly overlap at cell volumes of ~10^0^‒10^2^ μm^3^ (Fig. 2A). This overlap includes large bacteria with entirely active cytoplasms (e.g., *Azotobacter chroococcum*, *Magnetobacterium bavaricum*, ‘*Candidatus* Uab amorphum’, and *Chromatium okenii*; Dataset S1), picoeukaryotes which are relatively reduced (e.g., algae such as *Chaetoceros calcitrans*, *Micromonas pusilla*, *Nannochloris* sp., and *Nannochloropsis geditana*; Dataset S1), and phylogenetically diverse nanoeukaryotes (e.g., heterotrophic flagellates such as *Andalucia godoyi*, *Mantamonas plastica*, *Bodo saltans*, *Malawimonas jakobiformis*, *Palpitomonas bilix*, *Ancyromonas mylnikovi*, *Reclinomonas americana*; Dataset S1). Many small eukaryotes (both parasitic and free-living) can thus have sizes similar to those of many bacteria.

The histogram for genome size follows a similar pattern to that of cell volume: prokaryotes and eukaryotes have distinct but overlapping distributions (Fig. 2B). The genome size range for prokaryotes is <1‒16 Mbp, whereas that of eukaryotes is ~8‒10,000 Mbp. This suggests that there is an upper genome size constraint to prokaryotes based on the currently available data. Prokaryotes and eukaryotes also overlap at genomes sizes of ~8‒16 Mbp if the genomes of highly reduced eukaryotic parasites are excluded (Fig. 2B; Dataset S1). Many eukaryotes (protists) thus have genome sizes smaller than those of some prokaryotes. For example, prokaryotes such as myxobacteria, actinomycetes, cyanobacteria, and planctomycetes may have genomes of up to 16 Mbp in size (Dataset S1). The smallest genomes for free-living eukaryotes are those of some small green algae, red algae, and yeasts (8‒13 Mbp); some parasitic eukaryotes have genome sizes of just 2 or 6 Mbp (e.g., *Encephalitozoon* and *Babesia*; Dataset S1). The small heterotrophic nanoflagellate *Andalucia godoyi* (Jakobea), which have the most ancestral-like mitochondrial genomes, has a nuclear genome size of ~20 Mbp (26), barely larger than the largest prokaryotic genomes. For gene number, there is an even wider overlap between prokaryotes and eukaryotes (Fig. 2C). Prokaryotes with the greatest number of genes have 10,000‒13,000 genes (Dataset S1), whereas eukaryotes with the lowest number of genes include intracellular parasites (~2,000 genes in *Encephalitozoon*), free-living fungi (~4,500 genes in *Malassezia restricta* or ~6,400 in *Saccharomyces cerevisiae*), and small algae (~5,300 genes in *Cyanidioschyzon* and ~7,800 genes in *Ostreococcus tauri*). Some of the closest relatives of animals, the free-living flagellate *Monosiga brevicollis* (Choanoflagellata) and the symbiotic amoeba *Capsaspora owczarzaki* have ~9,200 and 8,800 genes, respectively (Dataset S1). In summary, the data suggest that, though there is some overlap between prokaryotes and eukaryotes, there also appears to be upper constraints to the cell volumes and genome size that prokaryotes can attain.

### The respiratory membrane requirement and maximum possible volume of cells

The cell volume of prokaryotes is potentially constrained by respiration at the cell surface (or by not having mitochondria; see introduction). In terms of energy, the rate of ATP synthesis at the cell surface must meet the rate of ATP consumption by the whole cell volume. However, surface area decreases relative to volume as cells grow larger—surface area scales with the square of length, whereas volume scales with the cube of length (27–29). A developmental or evolutionary increase in cell volume thus poses a challenge to cells because, if internal volumes remain active, processes that are carried out at the cell surface (e.g., respiration or nutrient transport) will, at some cell volume, be unable to support processes that occur in the cytoplasm (e.g., protein synthesis). Such scaling sets a maximum volume that cells cannot overcome in the absence of structural adaptations (e.g., mitochondria and endomembranes in eukaryotes, and intracytoplasmic membranes in prokaryotes (30)).

To determine the volumes at which cells first face a deficit in respiratory membrane, we examined the ratio between the amount of respiratory membrane needed and the maximum amount of respiratory membrane possible for a simple mitochondrion-less cell (i.e., a prokaryote). This ratio provides a measure of respiratory deficit, or the degree to which there is an excess or dearth of surface area allocated to respiration. Note that we do not assume any major structural adaptations (e.g., internalized membranes, external membrane protrusions or appendages, or internal inert spaces). The amount of respiratory membrane needed can be defined as the membrane area occupied by all respiratory complexes (or respiratory units) that are required to sustain the volume of a cell throughout its life cycle (*A*_*needed*_, in μm^2^). The maximum amount of respiratory membrane, in turn, is defined as the largest possible membrane area that can be devoted to respiratory complexes by a cell (*A*_*possible*_, in μm^2^). This area is necessarily only a fraction (*f*_*max*_) of the total membrane area available (*A*_*total*_, in μm^2^) because a cell also needs to allocate some of its surface area to lipids, nutrient transporters, protein translocases, flagella, etc.; *f*_*max*_ thus represents the maximum fraction of the total surface membrane that can be used for respiration. The respiratory deficit can then be expressed as:

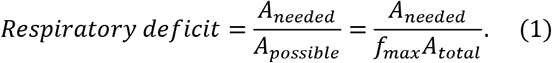

The amount of respiratory membrane needed by a cell can be calculated by multiplying the number of respiratory units (i.e., a complete set of respiratory complexes including the ATP synthase) required and the membrane area that each one of them takes up (*A*_*r*_). The number of respiratory units can be estimated by dividing the metabolic rate of a cell (*R*, in ATP h^−1^) by the ATP production rate of a single respiratory unit (*r*, in ATP h^−1^; Table S1). The metabolic rate of a cell can be expressed as the total ATP budget of a cell throughout its life cycle (*E*_*t*_) divided by the length of the cell cycle (or cell division time, *t*_*d*_ in h). The total energy budget of a cell (in ATPs) comprises both growth and maintenance costs (*c*_*g*_ and *c*_*m*_, respectively) and is calculated as in (10); this is adjusted to only include direct costs (*f*_*d*_) (31). The metabolic rate calculations agree with those reported previously by Chiyomaru and Takemoto, and are thus validated by empirical data (19) (Fig. S1). The amount of respiratory membrane needed by a cell thus depends on its energy demands, cell cycle length, and rate of ATP synthesis and area occupied by a single respiratory unit:

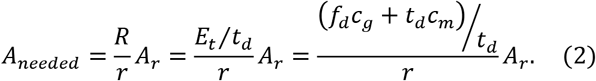

If the respiratory deficit is expressed as a function of cell volume (Supplementary Information), we obtain:

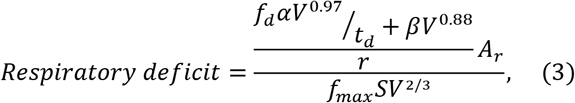

where *S* is a factor that specifies the shape of a cell (e.g., a perfect sphere or differently flattened spheroids; Supplementary Information). The parameters *f*_*d*_, *α*, *β*, *A*_*r*_, and *r* are constants whose values have been previously determined (10, 31–34) (Table S1). The parameters *t*_*d*_, *f*_*max*_, and *S* are constrained within biologically plausible ranges. For example, *t*_*d*_ is varied between 1 and 10 h, corresponding to the lower range of prokaryotic cell division times and the geometric mean of eukaryotic cell division times, respectively (10). The *f*_*max*_ parameter is varied between 8 and 18%, which are the largest possible fraction of respiratory membrane in *E. coli* (34) and the membrane fraction at which roughly half of all membrane proteins are respiratory enzymes (35). The shape factor, *S*, is varied between 4.8 and 12.1, which correspond to a sphere and an oblate spheroid with a cell length-width ratio of 0.1 (Fig. S2, Supplementary Information).

To assess the maximum possible volume that mitochondrion-less cells can achieve, we calculated the deficit in respiratory membrane (Eq. 3) across a wide range of cell volumes (Fig. 3A). Values of < 1 for the respiratory deficit indicate that the cell has an excess of respiratory membrane, whereas values of > 1 indicate that the cell has insufficient respiratory membrane to sustain its own volume. Respiratory deficit values of 1 thus point at the maximum volume that a simple mitochondrion-less cell can achieve, or the volume above which simple cells start to ‘gasp for air’. Our analyses show that spherical cells with a cell division time of 1 h and a maximum respiratory membrane fraction of 8% are surface area-constrained above a cell volume of about 1 μm^3^ (blue line; Fig. 3A). These parameter values and the estimated cell volume limit agrees with what is seen for a small and fast-growing bacterium like *Escherichia coli* (34). If half of membrane proteins are respiratory enzymes (i.e., a maximum respiratory membrane fraction of 18%), the largest volume that a cell can achieve is about 10 μm^3^ (black line; Fig. 3A). This large fraction of respiratory membrane would be possible if a bacterium devotes less of its surface area to other processes (e.g., flagella or chemotactic receptors), or alternatively, if a bacterium develops intracytoplasmic membranes for respiratory processes (30). A similar cell volume limit of 10 μm^3^ is achieved if the cell shape is changed to that of an oblate spheroid with a cell length-width ratio of 0.1 (dashed black line; Fig. 3A); some small and flattened flagellates like the eukaryote *Petalomonas minor* (36), or the phagocytic amoeboid prokaryote *Uab amorphum* have such cell body shapes (37). The cell volume limit is raised even more, to about 500 μm^3^, if the cell division time is increased to 10 h (dotted black line; Fig. 3A). The combination of these three changes raises the cell volume limit to higher than 10^5^ μm^3^ (red line; Fig. 3A). This might correspond to giant bacteria like *Thiomargarita* and *Epulopiscium* whose active cytoplasm is restricted to a thin enveloping sheet (i.e., 2% of the whole cell volume (38)), have long cell division times (1‒2 weeks (39)), and develop extensive intracytoplasmic membranes (e.g., *Epulopiscium* (40)).

**Figure 3.**
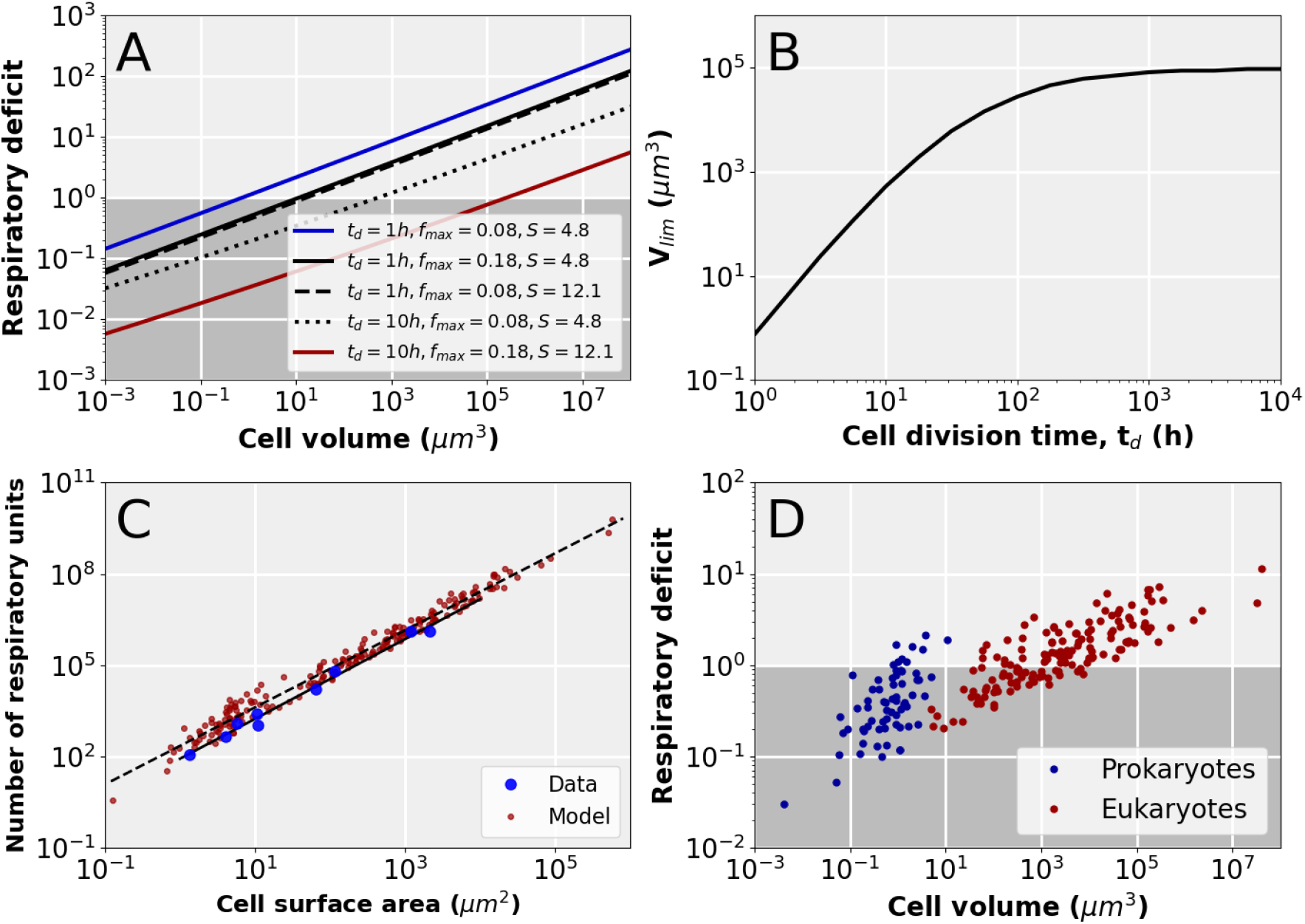
Factors that affect the volumes at which simple cells become constrained by their surface. **A**. The respiratory deficit as a function of cell volume. The blue line reflects cells that have a cell division time (*t*_*d*_) of 1 h, a maximum membrane occupancy of respiratory proteins (*f*_*max*_) of 8%, and a shape factor (*S*) of 4.8. The black lines reflect cells for which a single parameter, either *t*_*d*_, *f*_*max*_, or *S*, has been changed (see inset). The red line reflects cells for which all parameters have been simultaneously changed. The dark grey area indicates the domain in which there is enough surface area for respiration to support cell volumes. The intersection between each line (a defined set of parameters; see inset) and a respiratory deficit of one determines the maximum volumes that cells can achieve. **B**. The surface area-limited cell volume, *V*_*lim*_, plotted as a function of the cell division time. Here, fold deficit = 1, *f*_*max*_ = 8% and *S* = 4.8. **C**. The number of respiratory units (or ATP synthases) as a function of cell surface area. Empirically determined numbers of respiratory units (represented by ATP synthases) and cell surface areas, for prokaryotic and eukaryotic species, were obtained from (21) (blue points). The number of respiratory units was calculated (red points) using: ((*f*_*d*_*αV*^0.97^/*t*_*d*_) + *βV*^0.88^)/*r*, with the cell volumes and cell division times, for a range of prokaryotic and eukaryotic species, obtained from (10). The solid line is a fit to the data: y = 83 x^1.31^. The dashed line is a fit to the model: y = 221 ×^1.27^. **D**. Respiratory deficit calculated for individual prokaryotic and eukaryotic species whose cell volumes and cell division times have been previously estimated (10). Here, *f*_*max*_ = 8% and *S* = 4.8 (spherical cells).

Cells with longer cell cycles have lower metabolic rates and thus require fewer respiratory units (Eq. 2). This is because longer cell division times allow cells to accumulate the same amount of ATP required for growth over longer times spans. Thus, cells with longer cell division times can achieve larger cell volumes. Our model predicts that a spherical cell with a maximum respiratory membrane fraction of 8% can, potentially, reach an upper volume of about 10^5^ μm^3^ at a cell division time of roughly 10^3^ h (Fig. 3B). However, the cumulative amount of ATP required for cell maintenance continues to increase throughout the cell cycle (10), and this eventually limits the maximum cell volume that is possible (Fig. 3B).

Our model allows us to predict the number of respiratory units and amount of respiratory membrane area required by a cell (Eq. 3). The number of respiratory units predicted by our model follows closely, in both scaling exponent and intercept, the empirical data on the number of ATP synthases of cells previously reported by Lynch and Marinov (21) (Fig. 3C). Similarly, the amount of respiratory membrane required by eukaryotic cells also follows the data on mitochondrial inner membrane areas reported by Lynch and Marinov (21) after adjusting for the cross-sectional surface areas and stoichiometries of mitochondrial respiratory complexes (41) (Fig. S3).

We also calculated the respiratory deficit for prokaryotes and eukaryotes whose cell volumes and cell division times have been determined empirically (10). For these calculations, we assumed a spherical cell body shape (*S* = 4.8) and a maximum respiratory membrane fraction of 8%. These analyses showed that simple cells may have eukaryote-like volumes of up to 10^4^ μm^3^ without a shortage of surface area for respiration (dark area; Fig. 3D). Therefore, many eukaryotes might be able, theoretically, to support their cell volumes by respiring at their cytoplasmic membranes (i.e., without the need for internalizing respiratory membranes). Overall, our analyses thus reveal that longer cell cycles (*t*_*d*_), flattened or elongated cell shapes (*S*), and a larger allocation of surface area to respiration (*f*_*max*_) can, together or in isolation, allow cells to obtain larger volumes without the need for expanded respiratory membranes (e.g., mitochondria). On the other hand, increasingly larger, rounder and faster-dividing cells have higher respiratory deficits (i.e., larger than one) and are thus dependent on an excess of respiratory membranes that cannot be fully accommodated on their cytoplasmic membranes.

### The energetic investments in DNA of cells with contrasting genomic designs

Another claimed advantage of mitochondria is a drastic increase in ‘energy per gene’ due to the asymmetric genomic design (or ‘bioenergetic architecture’ *sensu* Lane and Martin) that they allow for in eukaryotes (3, 16–18). Eukaryotes have both a single nuclear genome, and numerous small and specialized mitochondrial genomes that scale with cell volume (i.e., genomic asymmetry; Fig. 4). Prokaryotes, in contrast, only have a single genome type whose copy number scales with cell volume (i.e., genomic symmetry; Fig. 4). Therefore, if a prokaryote were the size of an average eukaryote, the massive increase in gene number that accompanies polyploidy would keep its amount of energy per gene roughly equal to that of an average prokaryote despite having a much larger volume and energy available (3). On the other hand, according to Lane and Martin’s logic (3), eukaryotes have much more energy available per gene expressed as their nuclear genomes and gene numbers do not scale up with cell volume (3).

**Figure 4.**
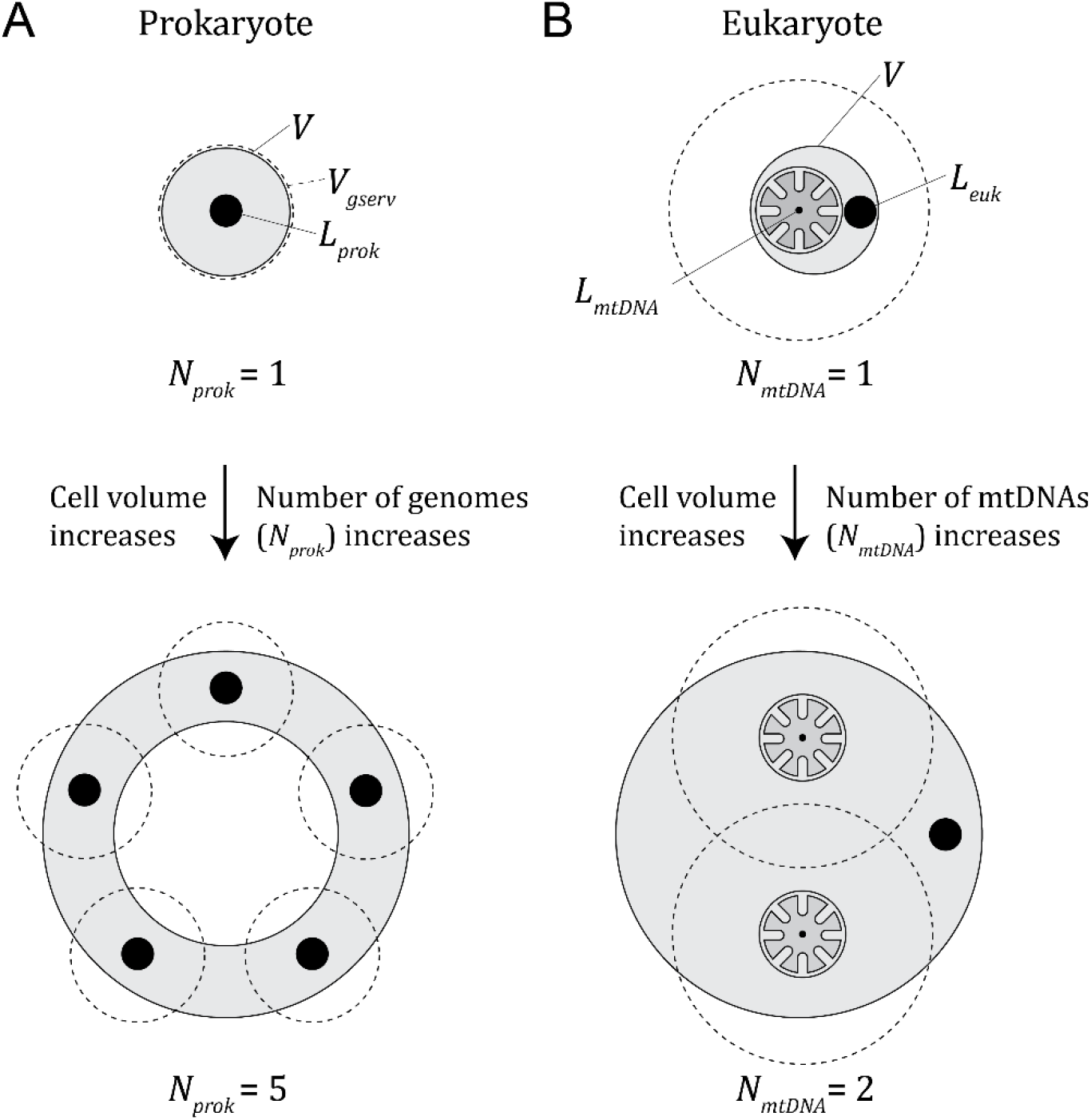
Graphical representation of contrasting genomic designs in prokaryotes and eukaryotes (Eq. 4 and see main text for an explanation of parameters). **A**. The genomic symmetry of prokaryotes. We have represented a large prokaryotic cell as a shell of cytoplasm surrounding a large inert central space, as seen in giant bacteria like *Epulopiscium* and *Thiomargarita*. Even though this cell architecture is irrelevant for our calculations (Eq. 5) as only the number of genomes is considered (filled black circles), prokaryotic cells have to scale up in cell volume with such an architecture remain viable in the absence of an active intracellular transport network (42). The total number of genomes *N*_*prok*_ is a function of the ratio of the cell volume and the volume controlled by a single genome (i.e., *V*/*V*_*gserv*_; see Eq. 5). **B**. The genomic asymmetry of eukaryotes. The dashed circles hypothetically represent the amount of volume that can be energetically supported by mitochondria. Because of cristae (expanded internalized respiratory membranes), mitochondria can, in principle, energetically support large cytoplasmic volumes. The total number of mitochondrial genomes *N*_*mtDNA*_ is a function of the total volume of mitochondria and the number of mtDNA molecules per μm^3^ of mitochondrial volume (*n*_*mtDNA*_*f*_*mt*_*V*; see Eq. 6 and main text) (43–45).

The concept of ‘energy per gene expressed’ has been criticized as having no evolutionary relevance (see (10, 22, 23)). This concept, as used by Lane and Martin, heavily penalizes large prokaryotes as their gene numbers increase with polyploidy. However, the amount of gene expression from each gene, irrespective of how many times the gene is duplicated, is proportional to cell volume. In other words, the relative cost of a gene, a more evolutionary meaningful concept (10, 21), remains constant. This is because the energetic demands of cells strictly depend on their cell volumes, i.e., prokaryotes and eukaryotes of the same volume require the same amount of energy. The concept of ‘energy per gene’ thus unfairly penalizes prokaryotes, or any polyploid. Furthermore, measurements of ‘energy per gene’ previously performed by Lane and Martin (3) unfairly favor eukaryotes because gene copies due to mitochondrial genome polyploidy (which scale with cell volume) were ignored (3). Because the concept of ‘energy per gene’ is inappropriate, our approach below thus relies on estimating the cost of cellular features (i.e., DNA synthesis) relative to the entire energy budget of a cell (10, 21).

To test the hypothesis that the genomic design of eukaryotic cells provides an overwhelming advantage, we developed an explicit model that compares the energetic capacity of eukaryotes to prokaryotes. The goal here is to isolate the genomic design of a cell from other confounding factors that also separate eukaryotes from prokaryotes. Because ATP demands depend on cell volume (and not complexity or gene number (10, 19)), we considered the amount of ATP that remains (1 − *C*_*DNA,euk*_ and 1 − *C*_*DNA,prok*_) after accounting for the relative cost of DNA that is associated with each genomic design (*C*_*DNA,euk*_ and *C*_*DNA,prok*_; Fig. 4 and Eq. 4). This remaining amount of ATP is devoted to all cell processes other than DNA synthesis (e.g., translation, transcription, lipid biosynthesis, etc.); the more ATP a cell invests in DNA, the less ATP there is to sustain other cellular processes. The ratio between the remaining ATP of a mitochondrion-bearing and mitochondrion-less cell thus provides a measure of the energetic advantage (>1), or disadvantage (<1), that mitochondrion-bearing cells might have. This can be expressed as:

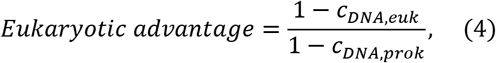

To calculate the cost of DNA for a prokaryote (a mitochondrion-less cell), we consider a cell with only a single main genome type. In prokaryotes, the number of genomes increases proportionally with cell volume, as seen in *Synechococcus elongatus* (46) or in giant bacteria like *Epulopiscium* (47) (see Fig. 4). The cause of this scaling might be the need to either bypass a diffusion constraint in the absence of active intracellular transport (42) or maintain genomes physically adjacent to respiratory membranes for efficient regulation (3, 18). We compiled data for several prokaryotes that show that the cell volume per genome does not exceed 2 μm^3^ in prokaryotes (*V*_*gserv*_ in Eq. 5; see Table S2). Our model thus assumes that if cell volume increases, the number of genomes must increase accordingly. The absolute total cost of DNA (in units of ATP) for a prokaryotic cell is the product of the amount of ATP required for synthesizing a single base pair (101 ATPs), the length of a single genome in base pairs (*L*_*prok*_ in Eq. 5) and the number of genomes. The number of genomes is the ratio between the total cell volume (*V*) and cell volume serviced by a single genome (*V*_*gserv*_) (Eq. 5). Finally, to obtain the relative cost of DNA for a prokaryotic cell, the absolute cost of the DNA is divided by the total ATP budget of the cell throughout its life cycle (Eq. 5). This can be expressed as a function of cell volume (10):

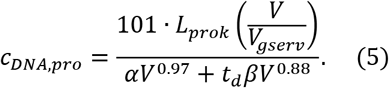

To calculate the cost of DNA for a eukaryote (a mitochondrion-bearing cell), we consider a cell with a single main (nuclear) genome and a variable number of mitochondrial genomes (mtDNA). If the cell volume increases, the number of mitochondrial genomes increase proportionally, but the main genome does not. The total number of mitochondrial genomes (*N*_*mtDNA*_ in Fig. 4) is the number of mtDNA molecules per μm^3^ of mitochondrial volume (*n*_*mtDNA*_) (43–45) multiplied by the total mitochondrial volume of the cell (*f*_*mt*_*V*) (Eq. 6). We compiled data that show that the cell volume fraction occupied by mitochondria (*f*_*mt*_) ranges from 1‒20% across diverse eukaryotes; our calculations thus use the geometric mean of 4.4% (Fig. 5A; see Table S3). The number of mtDNA molecules per nucleoid (or per μm^3^ of mitochondrial volume; *N*_*mtDNA*_), and the size of the mitochondrial genome (*L*_*mtDNA*_), were varied between 1-100 and 10 and 70 kbp (48, 49), respectively, with negligible effects on the results (Fig. S4). The total cost of DNA thus comprises the cost of the main genome and of all mitochondrial genomes required to support the whole cell volume (Eq. 6). The relative cost of DNA for a eukaryotic cell is calculated as above. If expressed as a function of cell volume, we have:

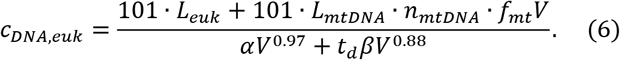

**Figure 5.**
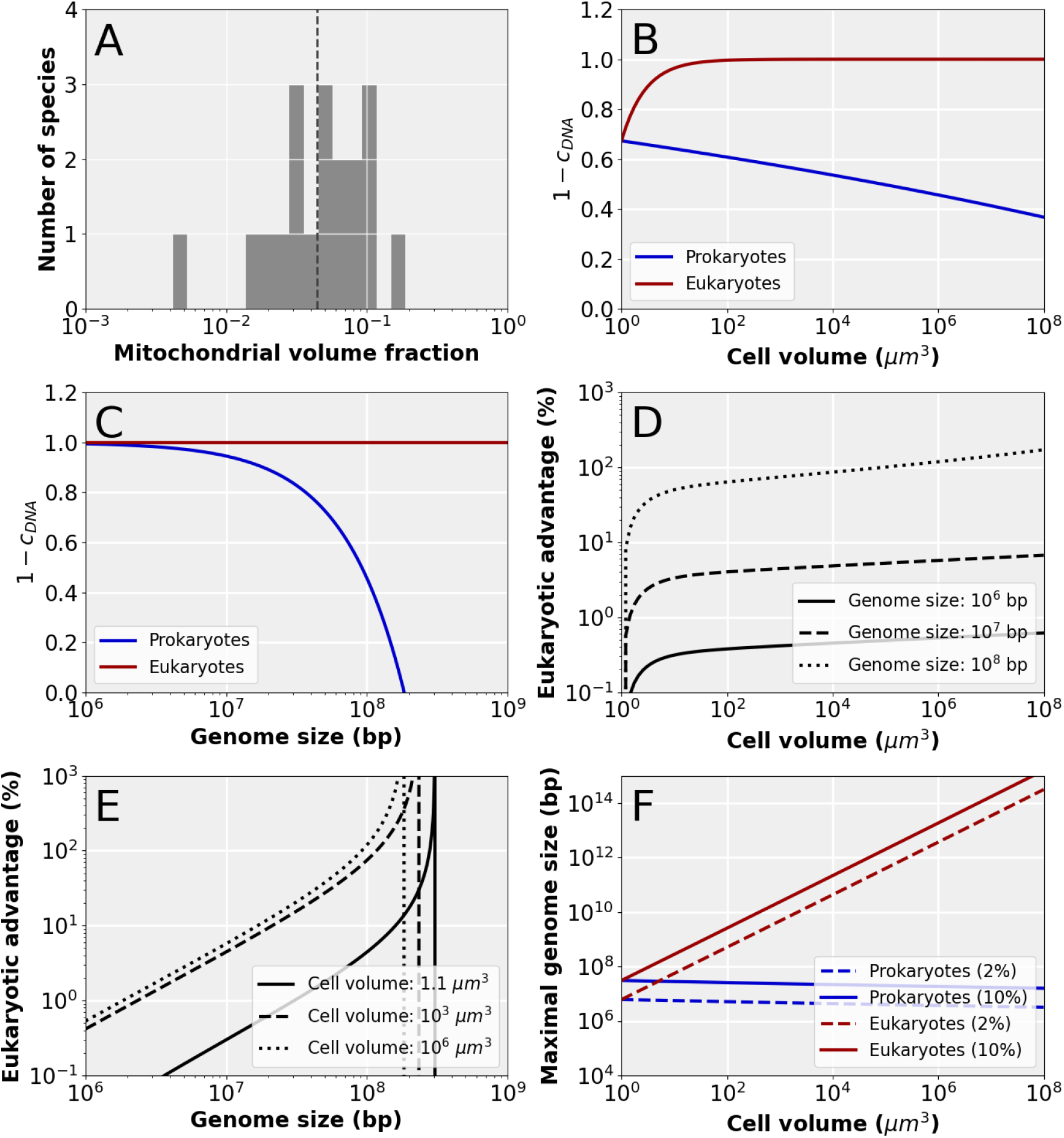
The impact of genomic design on energy allocation in cells. **A**. Distribution of mitochondrial volume fractions across a sample of phylogenetically diverse eukaryotes. The vertical dashed line indicates the geometric mean, 0.044 or 44%. **B**. The relative cell energy budget available for cellular features other than DNA as a function of cell volume. The plot was calculated with Eq. 5 and 6 and *L*_*euk*_ = *L*_*prok*_ = 10^8^ bp, *L*_*mtDNA*_ = 70 Kbp, *N*_*mtDNA*_= 10, *t*_*d*_ = 10 h, and *V*_*genome*_ = 1 μm^3^ (these values were also used for C-F). **C**. The relative cell energy budget available for cellular features other than DNA as a function of genome size. The plot was calculated with Eq. 5 and 6 and *V* = 10^6^ μm^3^. **D**. The energetic advantage of eukaryotes over prokaryotes as a function of cell volume. The plot was calculated with Eq. 4 and three different genome sizes as shown in inset. **E**. The energetic advantage of eukaryotes over prokaryotes as a function of genome size. The vertical lines denote the genome sizes at which the entire ATP budget of a prokaryote is devoted to DNA synthesis (1 − *C*_*DNA,prok*_ = 0). The plot was calculated with Eq. 4 and three different cell volumes as shown in inset. **F**. The maximum (main) genome size as a function of cell volume for prokaryotes and eukaryotes, for maximum DNA investments of 2 and 10 % of the entire cell energy budget. The plot was calculated with Eq. 5 and 6.

Our model (Eq. 4–6; see also Table S1) allows us to compare the contrasting genomic designs of eukaryotes and prokaryotes across a range of cell volumes and genome sizes (Fig. 5). Note that for these calculations, we kept the (main) genome size for eukaryotes equal to that of prokaryotes (i.e., *L*_*prok*_ = *L*_*euk*_) as we are only interested in determining whether the genomic asymmetry of eukaryotes provides an advantage over prokaryotes. We also kept the cell division time (*t*_*d*_) equal for prokaryotes and eukaryotes; varying *t*_*d*_ from 0‒100 h did not have major effects on the calculated eukaryotic advantage (Fig. S4), and thus does not change our conclusions. The main conclusions from our calculations are as follows. First, prokaryotes invest a larger fraction of their ATP budget on DNA as their cells increase in volume and, as a result, are left with less ATP for other processes such as gene expression (Fig. 5B). Second, the decrease in ATP available for other cell functions in prokaryotes is more pronounced as genome size increases (Fig. 5C). In contrast, eukaryotes suffer a negligible decline in their cellular ATP budget as their cell volume or main genome size increase (Fig. 5B, 5C). Third, eukaryotes have an energetic advantage (in terms of DNA cost savings) of less than ~120% for genome sizes of 10^6^−10^8^ bp and across a cell volume range of eight orders of magnitude or 10^0^−10^8^ μm^3^ (Fig. 5D; a volume of 10^0^ μm^3^ approximately corresponds to that of *Escherichia coli*, whereas a cell volume of 10^8^ μm^3^ is similar to that of a giant single-celled species like *Chaos carolinensis* (50)). At genome sizes of 10^6^−10^7^ bp, the energetic advantage of eukaryotes over prokaryotes is less than 10% across a similar range in cell volume (Fig. 5E). Fourth, a prokaryote with a genome size of 3×10^7^ bp, which is characteristic of many single-celled eukaryotes (see below), would have an energetic disadvantage of ~20% relative to a eukaryote with the same genome size. Such a genome size could, in principle, accommodate 2×10^4^ genes and up to ~1×10^7^ bp in regulatory sequences. Fifth, prokaryotic genomes cannot get larger than ~3×10^8^ because the cost of DNA would exceed the total ATP budget of the cell (at any cell volume). Eukaryotes, on the other hand, can achieve (main) genome sizes orders of magnitude larger as cell volume increases (Fig. 5F). If 2‒10% of the ATP budget of the cell is devoted to DNA synthesis, prokaryotes can reach genomes of 6×10^6^‒3×10^7^ bp in size (Fig. 5F).

## Discussion

The role of mitochondrial energetics in the origin and diversification of eukaryotes remains highly contested (3, 10, 21, 25, 51). As an attempt to resolve this debate, we investigated the respiratory deficit of mitochondrion-less cells and the maximum cell volume that can be supported by respiration at the cell surface. We showed that the maximum volume that a cell can attain is dependent on at least three major factors: cell body shape, cell division time, and maximum respiratory membrane fraction. A combination of biologically plausible values for these factors may allow mitochondrion-less cells to achieve volumes of up to 10^3^−10^5^ μm^3^ without a deficit in surface area devoted to respiration (Fig. 3). Furthermore, we investigated the energetic consequences of the contrasting genomic designs of mitochondrion-less and mitochondrion-bearing cells. Our results show that the asymmetrical genomic design of mitochondrion-bearing cells provide slight energetic savings in DNA costs relative to mitochondrion-less cells across a wide range of cell volumes and genome sizes (Fig. 5). The model further predicts that mitochondrion-less cells can achieve a genome size of 3×10^7^ bp, if they devote 10% of their ATP budget to DNA synthesis, at an energetic disadvantage of 20% (or 1.2-fold) (Fig. 5E, 5F).

The upper cell volumes and genome sizes of mitochondrion-less cells can be predicted based on energetic considerations, as done here. However, evolutionary success depends not only on the energetic capacity of a cell to sustain its own features, but also on the selective or ecological advantages conferred by such features. For example, a cell that has an energetic disadvantage by investing a large proportion of energy in DNA (and thus less in ribosome biogenesis or growth) but has a feature that confers a large ecological advantage (e.g., phagocytosis or antibiotic secretion) may otherwise outcompete cells that invest less in DNA but lack such a feature. Similarly, the reproductive disadvantage that may accompany longer cell division times in larger cells may be overcome by ecological specialization to avoid competition. This is the sentiment behind some of the criticisms of the energetic hypothesis for the origin of eukaryotic complexity previously raised by others (23, 24).

We have shown that the genomic design of eukaryotes may be advantageous in comparison to that of prokaryotes. However, this energetic advantage never exceeds 200% (or 3-fold) across a vast cell volume range of 10^1^‒10^8^ μm^3^ (Fig. 5B). These results stand in sharp contrast with the claim that ‘[an average] eukaryotic gene commands some 200,000-fold more energy than a prokaryotic gene’ (3). The discrepancy resides not only in the inappropriateness of the ‘energy per gene’ concept (see above), but also in that previous analyses compared idealized averages for modern eukaryotes and prokaryotes. Such averages, however, differ drastically in cell volumes. Because the energy demands of cells (i.e., ATP requirements and maximum metabolic rates (10, 19)) scale continuously and nearly linearly with volume, and prokaryotes and eukaryotes overlap across this continuum (Fig. 2A), such comparisons between rough averages are misleading.

The maximum advantage of 3-fold for eukaryotes found here stems from the comparison of two considerably different types of cells: mitochondrion-less and mitochondrion-bearing cells. Whereas the former represents an average prokaryote, the latter arguably represents a derived (proto-)eukaryote a with highly reduced mitochondrial genome and a dynamic cytoskeleton. This is because our model considers a mitochondrial genome that is less than 7% the size of the main genome (i.e., ≤70 Kbp, equivalent to that of jakobids and LECA), and assumes a main nuclear genome whose copy number does not increase with a larger cell volume. Such a reduced mitochondrial genome could only have evolved after the invention of a protein import machinery that sped up gene transfer to the nucleus or main genome (i.e., by allowing import of transferred gene products). In addition, only the presence of active intracellular transport (i.e., a dynamic cytoskeleton and motor proteins that bypassed diffusion constraints) would have allowed the nuclear or main genome *not* to scale up with cell volumes (unlike in prokaryotes (42, 47)). Thus, a great degree of evolutionary change (and time) separates the two types of cells compared here. This suggests, then, that the energetic advantages between immediate ancestor and descendant populations of proto-eukaryotes were necessarily much smaller than the 3-fold energetic advantage for mitochondrion-bearing cells found here. If the transition from FECA to LECA involved only 10,000 generations, and mitochondrial DNA deletions were uniformly distributed among them, the trans-generational selective advantage of an asymmetric genomic design was ~0.02%.

Comparative genomic analyses have estimated that the Last Eukaryote Common Ancestor (or LECA) had 4,431 gene domains (52), ~ 4,137–5,938 gene families (53–55) or 7,447–21,840 genes (mean = 12,753) (11). This inferred number of genes can be accommodated by a genome of ~20‒ 50 Mbp in size that also devotes more than a third of its size to regulatory and other non-coding DNA. Because nuclear DNA amounts scale strongly with cell volume in eukaryotes as the power law *V* = 1025.4 ∙ *DNA*^0.97^ (where *V* is in μm^3^ and *DNA* in pg) (56–58), a haploid LECA with such a genome size may have had a cell volume of ~23‒57 μm^3^ (i.e., the volume of a spherical cell of 3.5‒4.8 μm in diameter). Indeed, these genome and cell sizes are similar to those of small heterotrophic nanoflagellates such as jakobids and malawimonads which also have the most ancestral-like mitochondrial genomes known (48, 59). The gene number, genome size, and cell volume inferred for LECA fall within or is close to the modern prokaryote-eukaryote overlap (i.e., 10^0^‒10^2^ μm^3^, 10^6^‒ 10^7^ bp, and 4,000‒13,000 genes), and also encompass the cell volumes and genome sizes at which prokaryotes may not face a shortage of surface area (Fig. 3) or a considerable energetic disadvantage due to increasing DNA costs (Fig. 5). Thus, the prokaryote-eukaryote transition may have happened under these conditions.

Even though our analyses suggest that mitochondrion-less cells may achieve relatively large volumes and genome sizes under certain conditions, they also point at constraints that these simpler cells inevitably face at even larger volumes or genome sizes. Because the amount of respiratory membrane needed (i.e., number of ATP synthases) scales super-linearly with total surface area (21) (Fig. 3C and S3), prokaryotes may experience a shortage of respiratory membrane area at larger cell volumes (as long as their internal volumes remain active unlike in giant bacteria). Eukaryotes, on the other hand, can maintain such a super-linear scaling and reach much larger cell volumes by internalizing respiratory membranes in mitochondria. In other words, mitochondria allow energy supply to continuously match energy demand at increasingly larger volumes. Mitochondria may also allow eukaryotes to have shorter cell cycles and rounder (or less flattened) cell body shapes than mitochondrion-less cells (e.g., prokaryotes) at comparable cell volumes. Furthermore, as genome size increases, prokaryotes divest more and more of their ATP budgets to DNA synthesis, due to their genomic symmetry. Therefore, the energetic advantage of eukaryotes over prokaryotes increases with larger genome sizes. The maximum genome size that prokaryotes can theoretically achieve is 3×10^8^ bp if the entire ATP budget were devoted to DNA synthesis, or up to 3×10^7^ bp at 10% of the ATP budget. In contrast, eukaryotes can drastically expand their genomes as their cell volumes (and ATP budgets) grow larger, due to their genomic asymmetry. These theoretical predictions are consistent with constraints on prokaryotes suggested by the cell volume and genome size distributions (Fig. 2) and are at odds with the conclusions of Lynch and Marinov (10, 21).

## Conclusions

It has been claimed that an energy gap underlies the large differences in size and complexity between eukaryotic and prokaryotic cells. The proponents of this view further hold that the origin of mitochondria was a pre-requisite for simple prokaryotic cells to bridge such a gap and evolve into complex eukaryotic cells. Based on energetic considerations, we have shown here that prokaryotes can theoretically achieve eukaryote-like cell volumes and genome sizes. These findings are consistent with the modern prokaryote-eukaryote overlap in cell volumes and genome sizes. Because LECA was probably a small heterotrophic flagellate similar to a modern jakobid or malawimonad eukaryote, we suggest that the prokaryote-eukaryote transition did not necessarily require an expansion of respiratory membranes or the savings in DNA costs that mitochondria can provide. We also argue that the selective advantages conferred by mitochondria did not represent a quantum leap in energy supply (or ‘bioenergetic jump’ (3)) and were, in principle, not different from those provided by other eukaryotic innovations, such as a dynamic cytoskeleton or an endomembrane system. Mitochondria, however, became much more important for increasingly larger and faster-dividing eukaryotic cells, and may have thus allowed eukaryotes to successfully diversify and occupy novel adaptive zones throughout their evolutionary history.

## Acknowledgments

We thank Michael Lynch for comments on an early draft of this manuscript. SAM-G is supported by a EMBO Postdoctoral Fellowship (ALTF 21-2020). PES is supported by the Moore–Simons Project on the Origin of the Eukaryotic Cell, Simons Foundation 735927, https://doi.org/10.46714/735927.

## Notes

### Competing Interest Statement

The authors have declared no competing interest.

