## Supplementary Information for "The role of mitochondrial energetics in the origin and diversification of eukaryotes"

**This PDF file includes:**

Supplementary text
Figures S1 to S4
Tables S1 to S3
Legends for Datasets S1
SI References

**Other supplementary materials for this manuscript include the following:**

Datasets S1

**Supplementary Information Text**

**Derivation of Equation 3**. To determine the limits to cell volume we calculate the respiratory deficit, which we define as the membrane surface area needed for respiratory enzymes, $\boldsymbol{A_{needed}}$ (µm²), divided by the maximal surface area available to respiratory enzymes:

$$Respiratory\ deficit = \frac{A_{needed}}{f_{max}A_{total}}. \quad (S1)$$

Here, $A_{total}$ is the total cell surface area (in µm²), and $f_{max}$ is the maximal fraction of the cell surface area that can be devoted to respiratory enzymes. The value of $f_{max}$ must be below one because a cell membrane cannot function without lipids, and the cell cannot function without nutrient transporters, protein translocases, and other membrane proteins. The membrane surface area that is needed for respiration, $A_{needed}$, depends on the metabolic rate of the cell, $R$ (ATP h$^{-1}$), the ATP production rate of a respiratory unit, $r$ (ATP h$^{-1}$), and the area occupancy of a respiratory unit, $A_r$ (µm²):

$$A_{needed} = \frac{R}{r}A_r. \quad (S2)$$

A respiratory unit contains complex I-IV and ATP synthase in the stoichiometry reported by (1). The metabolic rate can be expressed as the total ATP budget, $E_t$, of the cell divided by the cell division time, $t_d$:

$$R = E_t/t_d. \quad (S3)$$

The total ATP budget, $E_t$, is a sum of two parts: (1) the amount of ATP needed to create a copy of a cell, and (2) the amount of ATP needed to maintain this cell. Following (2) and introducing a slight modification we obtain:

$$E_t = f_d c_g + t_d c_m, \quad (S4)$$

where $c_g$ is the growth cost (ATP), $c_m$ is the maintenance cost (ATP h$^{-1}$), and $t_d$ is the cell division time (h). We have introduced the factor, $f_d$, which is the fraction of the growth cost that is a direct cost. The direct cost counts the actual ATP molecules, whereas the other component of the growth cost, the opportunity cost, counts the potential ATP's that are sequestered in building blocks such as amino acids (3).

Combining Eq. S1-4 we obtain:

$$Respiratory\ deficit = \frac{\frac{{f_d c_g}/{t_d} + c_m}{r}A_r}{f_{max}A_t}. \quad (S5)$$

Next, we want to express the respiratory deficit as a function of cell volume. We do this by using empirically determined relations for $c_g$ and $c_m$ (2),

$$c_g = \alpha V^{0.97} \quad (S6)$$

$$c_m = \beta V^{0.88} \quad (S7)$$

and the mathematical relation between the surface area and volume of a 3D shape,

$$A_t = SV^{\frac{2}{3}}. \quad (S8)$$

Here, S is the shape factor, which has the value of $4\pi\left(\frac{3}{4\pi}\right)^{\frac{2}{3}} \approx 4.8$ for a sphere. Eq. S6-8 are substituted into Eq. S5 to obtain:

$$Respiratory\ deficit = \frac{\frac{f_d \alpha V^{0.97}/t_d + \beta V^{0.88}}{r} A_r}{f_{max} S V^{\frac{2}{3}}} = \frac{\left(f_d \alpha V^{0.97}/t_d + \beta V^{0.88}\right) A_r}{r f_{max} S V^{\frac{2}{3}}}, \quad (S9)$$

which is the respiratory deficit as a function of cell volume.

**Calculation of the shape factor, $S$.** The volume of a spheroid with semi-diameters of $a$, $a$, and $b$ is given by:

$$V_{sph} = \frac{4\pi}{3} a^2 b. \quad (S10)$$

The surface area is approximated by:

$$A_{sph} = 4\pi \sqrt[p]{\frac{a^{2p} + 2a^p b^p}{3}}, \quad (S11)$$

where $p$ = 1.6075.

The parameter $b$ can be expressed in terms of $a$ as:

$$b = ratio \times a. \quad (S12)$$

Thus, we can write:

$$V_{sph} = \frac{4\pi}{3} ratio \times a^3 \quad (S13)$$

and,

$$A_{sph} = 4\pi \sqrt[p]{\frac{(1 + 2ratio^p) a^{2p}}{3}}. \quad (S14)$$

Next, the semi-diameter $a$ is written in terms of volume:

$$a = \frac{3V_{sph}}{4\pi \times ratio}^{1/3}, \quad (S15)$$

and is substituted into Eq. S14 to obtain:

$$A_{sph} = 4\pi \sqrt[p]{\frac{(1 + 2ratio^p)\left(\frac{3V_{sph}}{4\pi \times ratio}\right)^{2p/3}}{3}} = 4\pi \sqrt[p]{\frac{(1 + 2 \times ratio^p)}{3}} \sqrt[p]{\left(\frac{3V_{sph}}{4\pi \times ratio}\right)^{2p/3}}$$

$$= 4\pi \sqrt[p]{\frac{(1 + 2 \times ratio^p)}{3}} \left(\frac{3V_{sph}}{4\pi \times ratio}\right)^{2/3}$$

$$= 4\pi \left(\frac{3}{4\pi}\right)^{2/3} \sqrt[p]{\frac{(1 + 2 \times ratio^p)}{3}} \left(\frac{1}{ratio}\right)^{2/3} V_{sph}{}^{2/3} = S V_{sph}{}^{2/3}. \quad (S16)$$

Thus,

$$S = 4\pi \left(\frac{3}{4\pi}\right)^{2/3} \sqrt[p]{\frac{(1 + 2 \times ratio^p)}{3}} \left(\frac{1}{ratio}\right)^{2/3}. \quad (S17)$$

For a sphere, $a = b$ and the $ratio$ = 1, which yields:

$$S = 4\pi \left(\frac{3}{4\pi}\right)^{2/3}. \quad (S18)$$

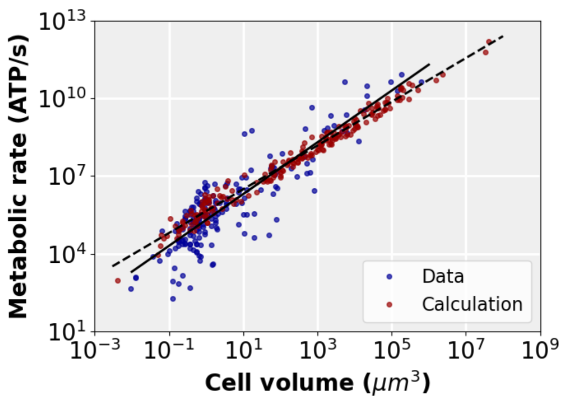


**Fig. S1. Comparing the results of metabolic rate calculations to data.** The blue points are empirically determined metabolic rates for various prokaryotic and eukaryotic species, obtained from (4) (units were converted by assuming that 1 mol ATP releases 50 kJ of energy). The red points are metabolic rates calculated with: $\boldsymbol{R = f_d \alpha V^{0.97}/t_d + \beta V^{0.88}}$, with the values for cell volumes and cell division times, for both prokaryotes and eukaryotes, obtained from (2). The solid line is a fit to the data: y = 2.0 × $10^5$ $x^1$, and the dashed line is a fit to the calculated points: y = 4.4 × $10^5$ $x^{0.85}$.

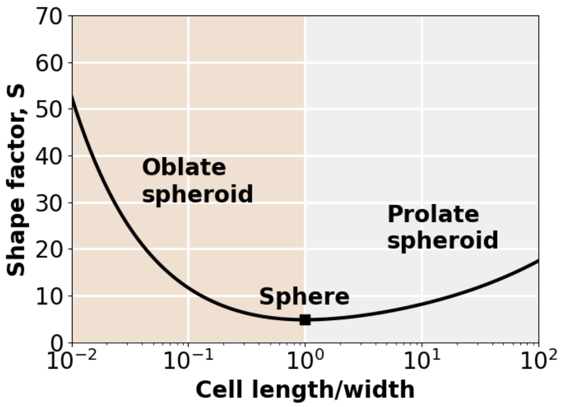


**Fig. S2: The shape factor, $S$, as a function of the ratio between cell length and width.** When this ratio is one, the cell is a sphere, and when this ratio is < 1 or >1, the cell is flattened into an oblate or prolate spheroid, respectively. The shape factor is calculated from Eq. S17.

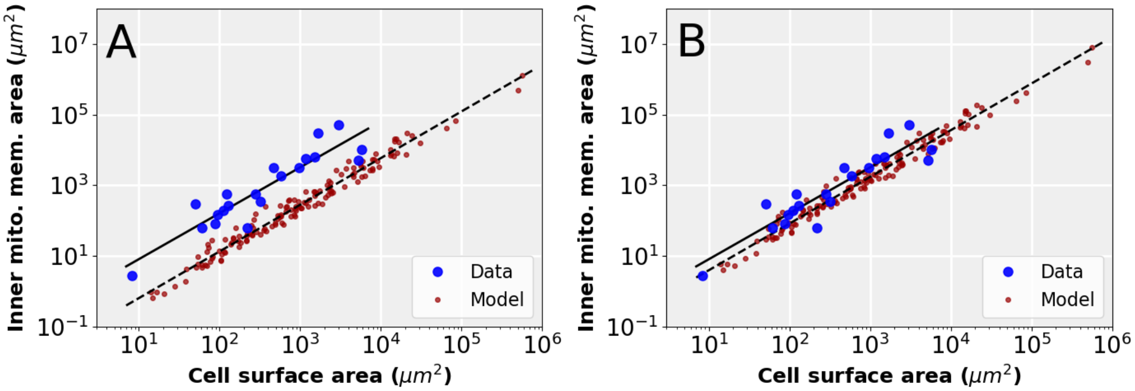


**Fig. S3: Prediction of the mitochondrial inner membrane surface area. A.** The inner mitochondrial surface area as a function of cell surface area. Empirically determined inner mitochondrial membrane areas were obtained from **(5)** (blue points). The inner mitochondrial membrane area was calculated (red points) with: $((\boldsymbol{f_d \alpha V^{0.97}/t_d}) + \boldsymbol{\beta V^{0.88}})/\boldsymbol{r} \cdot \boldsymbol{A_r} \times \mathbf{2.5}$, using cell volumes and cell division times for eukaryotic species obtained from **(2)**. The factor 2.5 was included to account for the lipids that support the membrane **(6)**. Note that for the calculation it is assumed that the inner mitochondrial membrane only houses respiratory proteins. The solid line is a fit to the data: y = 0.40 $x^{1.30}$. The dashed line is a fit to the model: y = 0.030 $x^{1.32}$. Here, the value of $\boldsymbol{A_r}$ is the one for *E. coli*, which is listed in Table S1. **B.** As for A except that the value of $\boldsymbol{A_r}$ used for the model calculations, which is dependent on both cross-sectional surface areas and stoichiometries of respiratory enzymes, is taken from a eukaryote (bovine) **(7)**, yielding a closer correspondence between data and model.

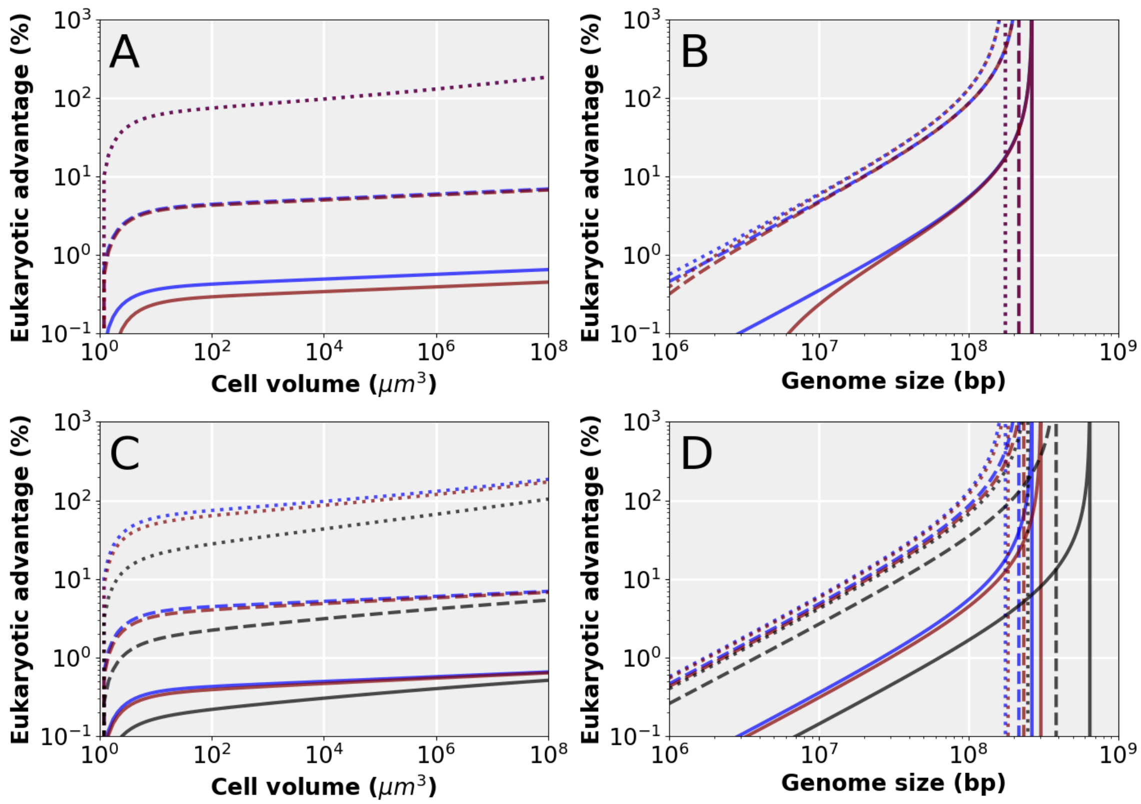


**Fig. S4. The effect of varying mitochondrial genome copy number, mitochondrial genome size, and cell division times on the eukaryotic advantage over prokaryotes.** Plots are generated from Eq. 4-6 with $V_{gserv}$ = 1 µm$^3$ and $f_{mt}$ = 0.044. **A, B**. Varying mitochondrial genome number and size. For the blue lines, $L_{mtDNA}$ = 10$^4$ bp and $n_{mtDNA}$ = 1 per µm$^3$ of mitochondrial volume. For the red lines $L_{mtDNA}$ = 7×10$^4$ bp and $n_{mtDNA}$ = 100 per µm$^3$ of mitochondrial volume. Cell division time, $t_d$ = 0. In some cases, red and blue overlap. **A**. For the dotted lines $L_{prok} = L_{euk}$ = 10$^8$, for the dashed lines $L_{prok} = L_{euk}$ = 10$^7$, and for the solid lines $L_{prok} = L_{euk}$ = 10$^6$. **B**. For the dotted lines $V$ = 10$^6$ µm$^3$, for the dashed lines $V$ = 10$^3$ µm$^3$, and for the solid lines $V$ = 1.1 µm$^3$. **C, D**. Varying cell division time, $t_d$. For all lines $L_{mtDNA}$ = 10$^4$ bp and $n_{mtDNA}$ = 1 per µm$^3$. For the blue lines $t_d$ = 0, for the red lines $t_d$ = 10 h, and for the black lines $t_d$ = 100 h. **C**. For the dotted lines $L_{prok} = L_{euk}$ = 10$^8$, for the dashed lines $L_{prok} = L_{euk}$ = 10$^7$, and for the solid lines $L_{prok} = L_{euk}$ = 10$^6$. **D**. For the dotted lines $V$ = 10$^6$ µm$^3$, for the dashed lines $V$ = 10$^3$ µm$^3$, and for the solid lines $V$ = 1.1 µm$^3$.

**Table S1. Parameter values for Eq. 3, 5, and 6.**

| Parameter (unit) | Value | References |
|---|---|---|
| $\boldsymbol{f_d}$ | 1/6 | (3) |
| $\boldsymbol{\alpha}$ (ATP) | $2.7\times10^{10}$ | (2) |
| $\boldsymbol{\beta}$ (ATP $h^{-1}$) | $3.9\times10^{8}$ | (2) |
| $\boldsymbol{A_r}$ ($\mu m^2$) | $8.4\times10^{-5}$ | (8, 1) |
| $\boldsymbol{r}$ (ATP $h^{-1}$) | $9.72\times10^{5}$ | (1, 9) |

**Table S2. Volume per genome for prokaryotes.**

| Species | Volume per genome ($\mu m^3$) | References |
|---|---|---|
| *Bacillus subtilis* | 0.66 | (10) |
| *Synechococcus elongatus* | 0.81 | (11) |
| *Buchnera* | 0.19 | (12) |
| *Methanosarcina acetivorans* | 1.1 | (13) |
| *Methanococcus maripaludis* | 0.03 | (13) |
| *Thermococcus kodakarensis* | 0.60 | (14) |
| *Epulopiscium fishelsoni* | 2.0 | (15) |

**Table S3. Mitochondrial volume fractions.**

| Species | Mitochondrial volume fraction | References |
|---|---:|:---:|
| *Ostreococcus tauri* | 0.077 | (16) |
| *Saccharomyces cerevisiae* | 0.014 | (17) |
| *Salpingoeca rosetta* | 0.059 | (18) |
| *Homo sapiens* (HeLa) | 0.071 | (19) |
| *Micromonas commoda* | 0.022 | (20) |
| *Pelagomonas calceolata* | 0.036 | (20) |
| *Emiliania huxleyi* | 0.051 | (20) |
| *Galdieria Sulphuraria* | 0.049 | (20) |
| *Phaeodactylum tricornutum* | 0.046 | (20) |
| *Symbiodinium pilosum* | 0.029 | (20) |
| *Mus musculus* | 0.085 | (21) |
| *Exophiala dermatitidis* | 0.099 | (22) |
| *Candida albicans* | 0.1 | (23) |
| *Tetrahymena pyriformis* | 0.15 | (24) |
| *Trichoderma viride* | 0.1 | (25) |
| *Chlorella fusca* | 0.03 | (26) |
| *Dunaliella salina* | 0.027 | (27) |
| *Medicago sativa* | 0.0046 | (28) |
| *Rhus toxicodendron* | 0.035 | (29) |

**Dataset S1 (separate file).** Dataset of cell volumes, genome sizes and gene number for a phylogenetically diverse set of prokaryotes and eukaryotes.
